# Systematic analysis of nonsense variants uncovers peptide release rate as a novel modifier of nonsense-mediated mRNA decay

**DOI:** 10.1101/2024.01.10.575080

**Authors:** Divya Kolakada, Rui Fu, Nikita Biziaev, Alexey Shuvalov, Mlana Lore, Amy E. Campbell, Michael A. Cortázar, Marcin P. Sajek, Jay R. Hesselberth, Neelanjan Mukherjee, Elena Alkalaeva, Zeynep H. Coban Akdemir, Sujatha Jagannathan

## Abstract

Nonsense variants underlie many genetic diseases. The phenotypic impact of nonsense variants is determined by nonsense-mediated mRNA decay (NMD), which degrades transcripts with premature termination codons (PTCs). Despite its clinical importance, the factors controlling transcript-specific and context-dependent variation in NMD activity remain poorly understood. Through analysis of human genetic datasets, we discovered that the amino acid preceding the PTC strongly influences NMD activity. Notably, glycine codons promote robust NMD efficiency and show striking enrichment before PTCs but depletion before normal termination codons (NTCs). This glycine-PTC enrichment is particularly pronounced in genes tolerant to loss-of-function variants, suggesting evolutionary selection or neutrality conferred by efficient elimination of truncated proteins from non-essential genes. Using biochemical assays and massively parallel reporter analysis, we demonstrated that the peptide release rate during translation termination varies substantially with the identity of the preceding amino acid and serves as the primary determinant of NMD activity. We propose a “window of opportunity” model where translation termination kinetics modulate NMD efficiency. By revealing how sequence context shapes NMD activity through translation termination dynamics, our findings provide a mechanistic framework for improved clinical interpretation of nonsense variants.

## INTRODUCTION

Proper interpretation of genetic variation is crucial for understanding and treating human disease. This challenge is particularly acute for nonsense variants, which account for over 30% of inherited human diseases due to the action of nonsense-mediated RNA decay (NMD) (Miller & Pearce, 2014). NMD eliminates transcripts that contain premature termination codons (PTCs) that would otherwise generate truncated protein products. Much of our understanding of NMD comes from studies of naturally occurring nonsense mutations and genotype-phenotype correlations in genes involved in Mendelian disorders such as β-globin (Neu-Yilik et al., 2011) (*HBB*), Dystrophin (Kerr et al., 2001) (*DMD*), and Lamin A/C (van Engelen et al., 2005) (*LMNA*). For example, individuals with a nonsense mutation in early exons of *HBB* develop a recessive form of β-thalassemia due to degradation of the mutant mRNA via NMD (Chang & Kan, 1979), whereas nonsense mutations in the last exon of *HBB* cause a dominant form of the disease due to the mutant mRNA escaping NMD and leading to the production of a C-terminally truncated β-globin protein (Kazazian et al., 1992). Such studies have helped refine the rules of NMD, including the “50-55 nt rule”, which describes the NMD insensitivity of nonsense mutations in the last and penultimate exons up to 55 nucleotides upstream of the final exon junction complex (Popp & Maquat, 2016).

Several exceptions to these rules have also been observed: nonsense mutations found close to the start codon often escape NMD due to downstream reinitiation of translation (Inacio et al., 2004; Pereira et al., 2015; Zhang & Maquat, 1997); stop codon readthrough has been reported to contribute to the escape of some nonsense variants from NMD (Hogg & Goff, 2010; Keeling et al., 2004); and NMD efficiency has been found to vary across cell types (Linde et al., 2007), tissues (Bateman et al., 2003; Zetoune et al., 2008), and individuals (Kerr et al., 2001; Nguyen et al., 2014), offering potential explanations for tissue tropism and varying penetrance across the population of disease-causing nonsense variants. Recent large-scale genomic studies have highlighted the limitations of our current predictive frameworks. A systematic study of PTCs in the Cancer Genome Atlas database found that the 55-nt rule only predicted the fate of ∼50% of PTCs; while additional factors such as exon length could account for another ∼20% of NMD variability, ∼30% of NMD variability is still unexplained (Lindeboom et al., 2016; Lindeboom et al., 2019). Another study investigating nonsense variants that are highly frequent in the healthy human population found that ∼50% of PTC-containing transcripts escape NMD via mechanisms including alternative splicing, stop codon readthrough, and alternative translation initiation (Jagannathan & Bradley, 2016). If every premature termination event that satisfies the 50-55 nt rule does not lead to NMD, what other factors influence the probability and activity of NMD?

As premature translation termination is the key signal for NMD, termination efficiency may impact NMD. Translation termination efficiency can be conceptualized in two ways: (1) the fidelity of termination, reflecting the likelihood of stop codon readthrough, and (2) the kinetics of termination, reflecting the dwell time of the terminating ribosome at the stop codon. Both of these aspects are influenced by cis-acting elements (Baker & Hogg, 2017; Loughran et al., 2014; McCaughan et al., 1995; Pierson et al., 2016; Tork et al., 2004). For example, specific elements within the stop codon sequence context and certain viral RNA structures can reduce termination fidelity (Baker & Hogg, 2017; Bonetti et al., 1995; Cridge et al., 2018; Loughran et al., 2014; Mangkalaphiban et al., 2021; Tork et al., 2004). Additionally, the identity of the nucleotide immediately after the stop codon and the peptide-tRNA hydrolysis rates of the amino acid preceding the stop codon impact the rate of peptide release (McCaughan et al., 1995; Pierson et al., 2016). These observations raise the possibility that local sequence context could serve as a key modulator of NMD activity, potentially explaining the observed variability in nonsense variant outcomes.

In this study, we hypothesized that the sequence contexts surrounding PTCs may hold the key to understanding NMD variability among transcripts. Indeed, we found that glycine codons are highly enriched immediately upstream of PTCs among healthy individuals, especially in nonessential genes, and that this sequence context potently enhances NMD activity. To systematically assess the impact of PTC sequence context on NMD activity, we performed a massively parallel reporter assay (MPRA) that enables comprehensive evaluation of sequence variants where NMD activity is calculated via RNA-seq readout of reporter transcripts with and without NMD inhibition, and found a broad range of NMD activities associated with different sequences. Statistical modeling of the MPRA data identified peptide release rate during translation termination as the major predictor of NMD activity, which we verified using biochemical assays. These findings reveal a previously unappreciated mechanism controlling NMD efficiency, where variable peptide release rates alter NMD activity by modulating the kinetic window available for NMD factors to bind and enact NMD during translation termination. Our data suggest that the Gly-PTC context, by virtue of allowing efficient elimination of nonsense-containing transcripts from non-essential genes, has risen to a higher frequency among healthy individuals through the course of evolution. Our results provide a new framework for predicting NMD efficiency that will enhance clinical interpretation of nonsense variants and guide therapeutic strategies.

## RESULTS

### Glycine codons are enriched preceding PTCs in non-essential genes

Healthy individuals harbor ∼100 putative loss of function genetic variants, most of which are protein-truncating (MacArthur et al., 2012). In previous work, we showed that such PTCs often escape NMD via mechanisms including alternative splicing or stop codon readthrough, allowing them to rise to high frequencies in the healthy population (minor allele frequency, MAF > 5%) (Jagannathan & Bradley, 2016). In this study, we focused on rare variants with MAF ≤ 1% in healthy individuals, with the assumption that such variants may be more likely to undergo NMD and yet have varying efficiencies. Using the gnomAD database (n=151,332, v2.1.1), which contains genetic variant data for >700,000 healthy humans (Karczewski et al., 2020), we analyzed amino acid enrichment within the 10 amino acids preceding PTCs (**Fig. 1A, Supp. Fig. 1A)**. We compared the enrichment of these amino acids to that of normal termination codon (NTC) contexts in the reference genome (n=23,066, **Fig. 1A, Supp. Fig. 1B**) using a method similar to Koren *et al* (Koren et al., 2018). Gly codons at the -1 position stood out among all examined positions as the most enriched before PTCs (**Fig. 1B-D**, and **Supp. Fig. 1**). As shown previously, Gly codons were least enriched before the NTC, consistent with their role as a C-terminal degron (Koren et al., 2018; Lin et al., 2018). This striking opposite trend for Gly enrichment upstream of a PTC versus an NTC suggests that it may be functionally relevant.

**Figure 1.**
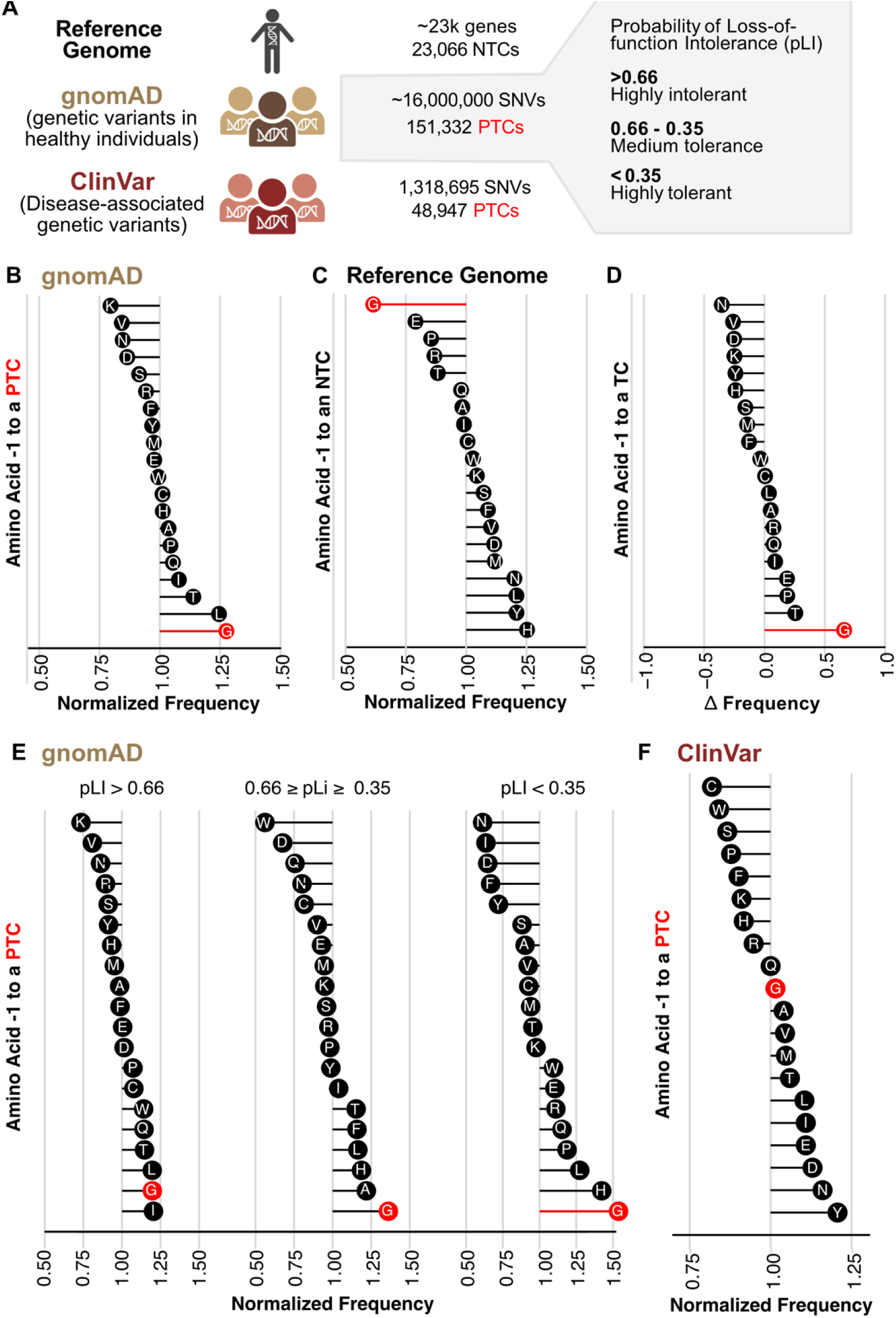
Gly is enriched preceding a PTC in non-essential genes. (**A**) Schematic for analyzing LoF scores for rare PTCs in healthy individuals and PTCs in disease contexts. (**B-C**) Amino acid enrichment - 1 to a PTC (B) or an NTC (C). (**D**) Each amino acid’s frequency delta (enrichment before a PTC-NTC) at the -1 position. (**E**) The -1 amino acid enrichment of PTCs binned into 3 categories of probability of loss-of-function intolerance (pLI). The lower the pLI, the more tolerant the PTCs are to loss of function. (**F**) The -1 amino acid enrichment of PTCs in disease contexts.

To further explore the functional impact of Gly-PTC enrichment in healthy individuals, we analyzed the relative frequency of PTC-inducing variants binned in terms of the probability of loss of function intolerance (pLI) score of their host genes (**Fig. 1A** and **1E**). pLI is determined by taking a gene’s estimated mutation rate and comparing how many protein-truncating variants (PTVs) are actually observed in a population versus how many would theoretically occur if these variants had no impact on fitness (Lek et al., 2016). The lower the pLI, the more tolerant to loss of function a gene is and the less essential. We found that Gly-PTC contexts were enriched in genes with lower pLI (**Fig. 1E**). We also examined the amino acids enriched immediately upstream of PTCs from the ClinVar database, which contains human genetic variants associated with disease (n=48,847, **Fig. 1A**) (Landrum et al., 2018). Intriguingly, we found no enrichment of Gly at the -1 position of disease-associated PTCs (**Fig. 1F**). In summary, the Gly-PTC enrichment in non-essential genes in comparison to essential or disease-relevant genes suggests a non-pathogenic and homeostatic role for this sequence context.

### Allele specific expression analysis shows Gly-PTC contexts lead to more efficient NMD

To explore the relationship between PTC sequence context and NMD efficiency, we calculated allele specific expression values using whole genome sequencing and RNA-seq datasets from the NHLBI Trans-Omics for Precision Medicine (TOPMed) program. For each stop gain variant (i.e. a single nucleotide variant that converts a codon to a premature stop) found in the TOPMed dataset, we measured allele-specific expression (ASE). ASE refers to the differing expression levels of alleles within a gene in a diploid organism, where, for example, a nonsense allele is less expressed compared to the “reference” or “wild type” allele due to NMD. A potential confounding factor in our analysis is the presence of other sources of allele-specific expression not associated with NMD. To minimize the effect of those factors, we filtered out variants in: 1) imprinted genes and genes with established cis-regulatory variants such as eQTLs (Baran et al., 2015; Consortium, 2013, 2015, 2020); 2) transcripts with stochastic low expression (<= 1 transcript per million (TPM)); and (3) single-exon genes that are not likely to be surveilled by EJC-enhanced NMD. Because the TOPMed program includes individuals with complex diseases and various experimental designs (cohort, case-control, etc.), we selected variants in genes highly tolerant to loss of function (pLI < 0.35) and excluded ultrarare variants (minor allele frequency, MAF <= 0.01%). This resulted in a final set of 883 heterozygous stop gain variants (**Fig. 2A**).

**Figure 2.**
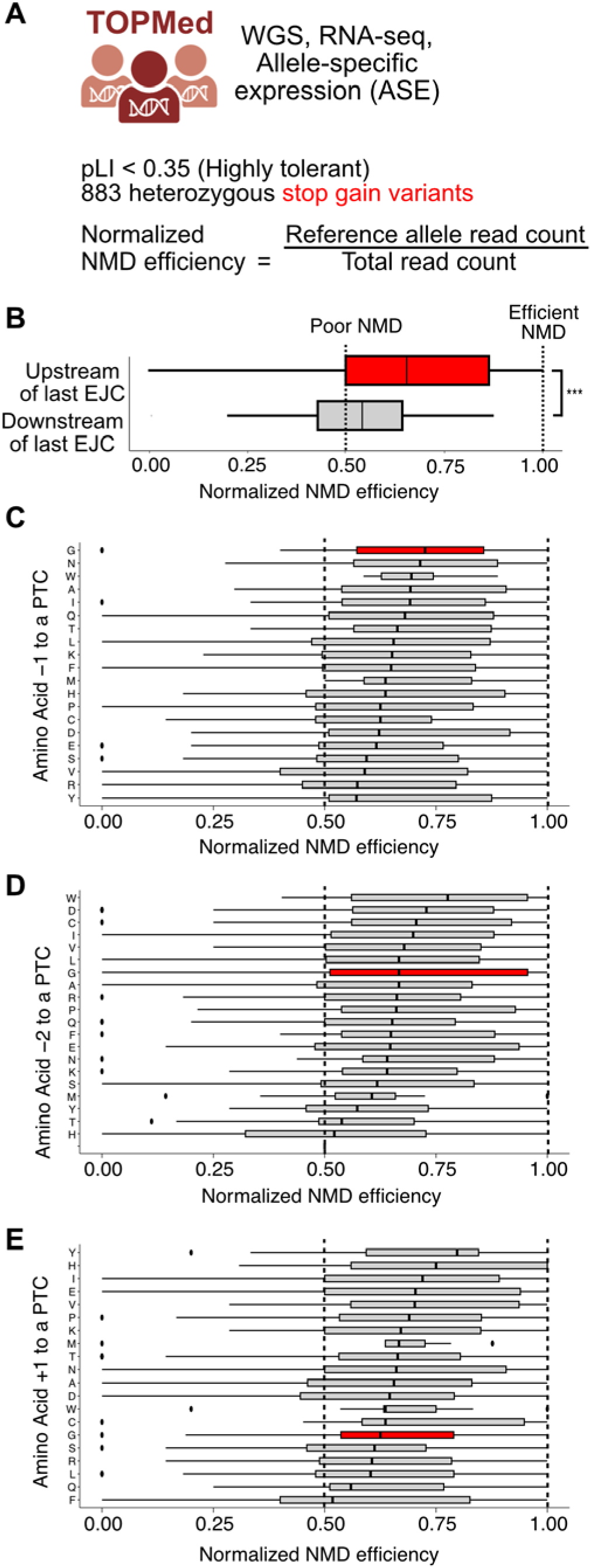
Gly at the -1 position leads to enhanced NMD efficiency. (**A**) Schematic for analyzing allele specific expression of heterozygous nonsense variants in TOPMed dataset. (**B**) A comparison of normalized NMD efficiency of variants with the PTC present upstream vs. downstream of the last EJC. Variants with the PTC upstream of the last EJC are NMD targets. (**C-E**) Normalized NMD efficiencies of nonsense variants categorized based on amino acids present at the -1 (C), -2 (D), and +1 (E) positions of the PTC.

As a positive control, we first confirmed that PTCs upstream of the last EJC triggered more efficient NMD compared to PTCs downstream of the last EJC (**Fig. 2B**), as would be expected based on the canonical 50-55 nt rule of NMD (Popp & Maquat, 2016). Next, we ordered stop gain variants according to the identity of their preceding amino acid codons and found that the variants with the amino acid glycine at the -1 position to the PTC showed the highest NMD efficiency (**Fig. 2C**). A similar trend was not observed for positions -2 and +1 (**Fig. 2D-E**). Together, these results suggest that having a glycine codon immediately upstream of a PTC leads to higher NMD efficiency compared to other sequence contexts, supporting a role for -1 codon identity in modulating NMD efficiency.

### Massively parallel reporter assay (MPRA) captures sequence-dependent variation in NMD activity

To systematically examine the relationship between NMD activity and the PTC sequence context, we conducted an MPRA using a set of NMD reporters we previously developed (Kolakada et al., 2024) (**Fig. 3A**). These reporters contain two types of NMD targets: one triggered by the presence of exon-junction complexes downstream of a PTC ("EJC-enhanced”) and the other by the presence of a long 3’ UTR (“EJC-independent”). Because these targets rely on related but distinct cues to trigger NMD, they might respond differently to PTC sequence variation.

**Figure 3.**
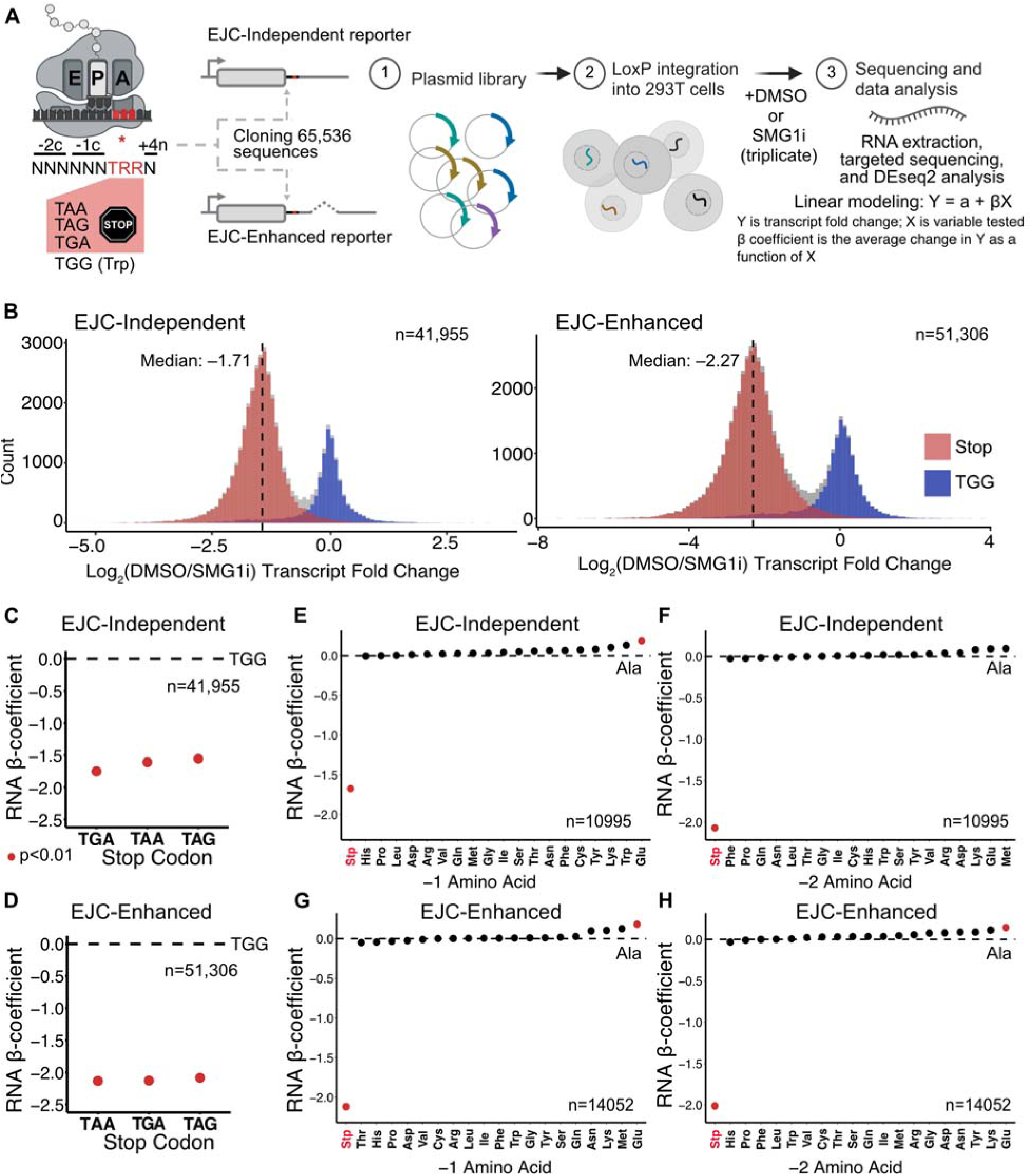
MPRA shows a wide range of PTC context-dependent NMD efficiencies. **(A)** Design of the MPRA. **(B)** Histogram of transcript fold change EJC-independent and EJC-enhanced NMD+ reporters comparing DMSO-treated cells to SMG1i-treated cells. Black dotted line represents the median of the stop codon population. **(C-D)** Dot plots of the β coefficient representing the effect of the stop codon on EJC-independent and EJC-enhanced NMD activity. β coefficients of all stop codons are in comparison to TGG **(E-H).** Plots depicting a variable’s effect on transcript fold change of TGG context reporters, β coefficients of all amino acids are in comparison to Ala: (**E-F**) Dot plot for the -1 (E) or -2 (F) amino acid for EJC-Independent reporters. **(G-H)** Same plots for EJC-enhanced reporters. All experiments were conducted in triplicates. All β coefficients with a p-value < 0.01 are plotted in red.

We varied the 6 nucleotides before the PTC, the PTC itself, and 1 nucleotide after the PTC in an unbiased fashion (N_6_TRRN; corresponding to 65,536 unique sequences; **Fig. 3A**) and asked how these sequences impacted NMD activity in both EJC-independent and EJC-enhanced reporters. Because we employed stop codons with “TRR” for array-based oligonucleotide synthesis (“R” being a purine), 25% of our sequence library consists of a tryptophan (Trp) codon (TGG) instead of a PTC – thereby serving as control sequences that do not undergo NMD. All sequences were incorporated into both EJC-independent and EJC-enhanced fluorescent reporter backbones (**Supp. Fig. 2A-B**) and integrated into a single genomic location in HEK293T cells via Cre-Lox mediated recombination (Khandelia et al., 2011). These cells were then treated with either DMSO or SMG1i, harvested for RNA, and subjected to targeted sequencing to quantify the levels of each sequence context in the pool of cells. The NMD activity of an individual reporter sequence was calculated as the log_2_ transcript fold change between DMSO-treated cells (NMD active) and SMG1i-treated cells (NMD-inhibited) as measured by DESeq2 (Love et al., 2014), thus a lower (negative) value indicates higher NMD activity, and vice versa. The ∼1/4^th^ of control sequences containing the Trp codon rather than a stop codon were used for normalization.

The NMD activity of both EJC-independent and EJC-enhanced reporters had a bimodal distribution with clear separation between constructs containing stop codons and the Trp-encoding TGG codon (**Fig. 3B**): sequences containing a stop codon were left-shifted compared to the sequences containing the Trp control, indicating that the former are targets of NMD. In the population of sequences undergoing NMD (i.e. those containing a stop codon), there was a wide range of NMD activity for both the EJC-independent and the EJC-enhanced reporters. EJC-enhanced reporter expression was reduced by 4.8-fold compared to the cells treated with the NMD inhibitor (median log2FC: -2.27), while the EJC-independent reporters experienced a 3.3-fold decrease (median log2FC: -1.71) (**Fig. 3B**). These results indicate that EJC-enhanced targets undergo more efficient NMD compared to EJC-independent targets, consistent with prior research (Buhler et al., 2006; Chu et al., 2021; Singh et al., 2008).

Next, to investigate whether different sequence elements explained the wide range of observed NMD activity (i.e. transcript fold change), we employed linear modeling. These models allowed us to evaluate the effect size (β coefficient) and statistical significance (p-value) of individual features within a variable on NMD activity relative to a control feature. For example, to understand the effect of stop codon identity on NMD activity, we evaluated the effect of the individual stop codons TAA, TAG, or TGA relative to TGG (Trp). We also evaluated the effects of the amino acids and the codons at the -1 and -2 positions and the nucleotide at the +4 position of the PTC on NMD activity. Negative β coefficients indicated stronger NMD activity, while positive ones indicated weaker NMD activity. It is important to note that each model (represented by individual plots) is unique, and the β coefficients cannot be quantitatively compared across the plots.

As expected, the linear models for the effect of stop codon identity on NMD activity showed that all stop codons caused significantly stronger NMD compared to the Trp (TGG) control for both EJC-independent and EJC-enhanced reporters (**Fig. 3C-D**). There were minor variations to the extent each stop codon affected NMD with TAG having the weakest NMD activity (highest β coefficient) across NMD targets (**Supp. Fig. 2C**). Notably, the TGA stop codon had a β coefficient 0.2 lower than TAG, indicating a 20% increase in NMD activity compared to the TAG stop codon for the EJC-independent reporters (**Supp. Fig. 2C**).

Before examining how other sequence elements such as the amino acids in the -1 and - 2 positions to the PTC affected NMD, we wanted to test whether these sequence elements affected the control TGG sequences. Linear models of TGG transcripts across NMD targets showed that the amino acids in the -1 and -2 positions had almost no significant effect on transcript fold change (i.e. transcript level in the DMSO condition normalized to SMG1i treatment) **(Fig. 3 E-H**). The exception is Glu, which caused a small but significant increase in transcript fold change at the -1 position for EJC-independent transcripts and both the -1 and -2 positions for EJC-enhanced transcripts (**Fig. 3 E, G-H**).

### PTC sequence context impacts EJC-independent NMD efficiency

We next constructed linear models to examine the effects of the amino acids and codons at the -1 and -2 positions before the PTC on NMD. The identity of the amino acid at the -2 position had a wide range of effect on EJC-independent NMD (**Fig. 4A**). As the -2 position is coded by randomized nucleotides (NNN), it can include all three stop codons. We found that a stop codon at the -2 position caused the strongest increase (∼40%) in NMD activity compared to the Ala control (**Fig. 4A**). More subtle effects were observed across the other amino acids, with Arg weakening NMD by ∼20% compared to Ala. When examining the effect of individual codons on NMD in a separate linear model, we found that the effects largely grouped by the identity of the encoded amino acid, with only a few that were significantly different (**Fig. 4B**; all codons compared to Ala-GCA)

**Figure 4.**
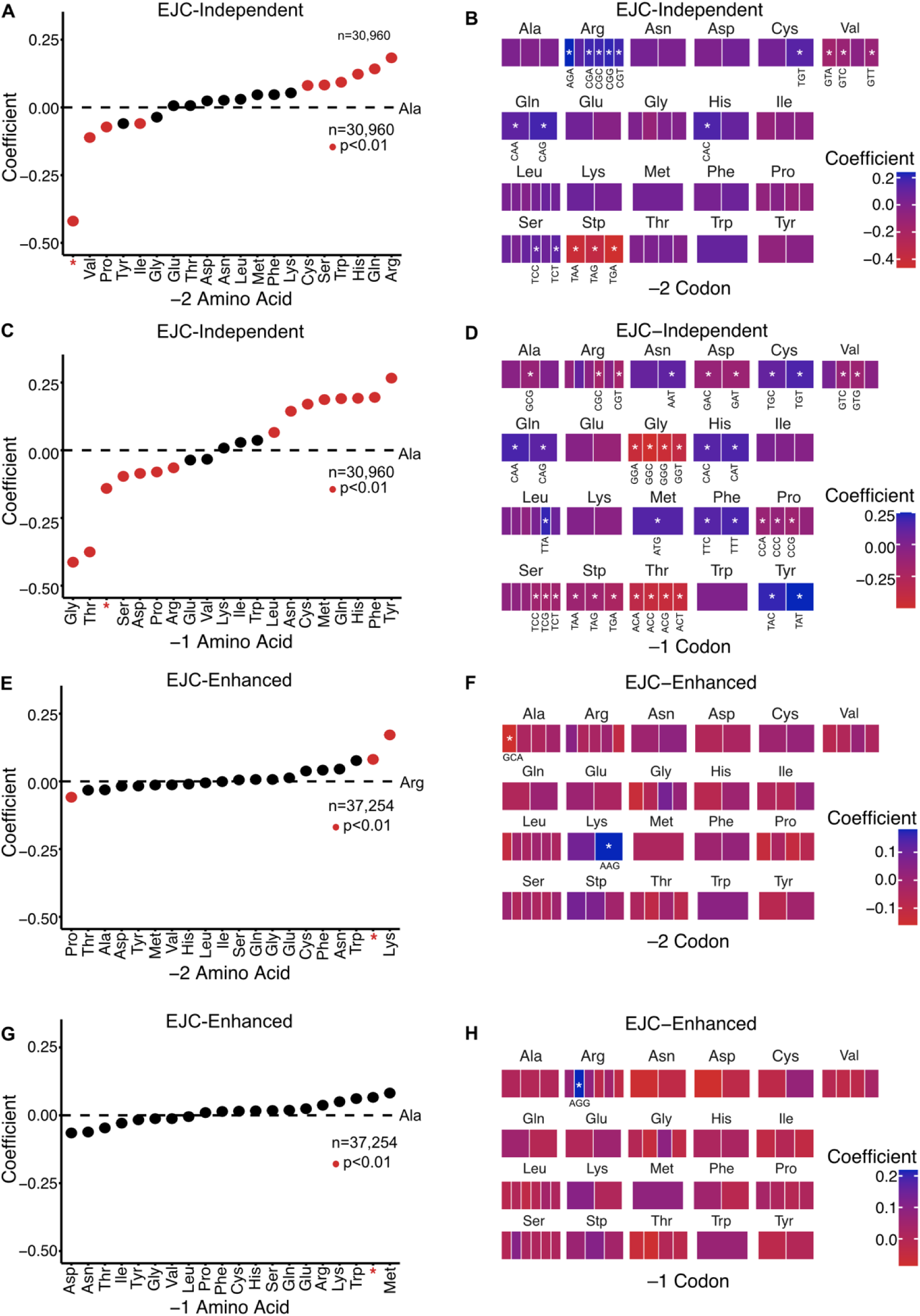
Effect of PTC sequence context on EJC-independent and EJC–enhanced NMD efficiencies. (A-H) Plots depicting the β coefficients for variable’s effect on EJC-independent **(A-D)** and EJC-enhanced **(E-H)** NMD activity: (A, E) Dot plot for the -2 amino acid and (B, F) Heat map for the -2 codon (C, G) Dot plot for the -1 amino acid, and (D, H) Heat map for the -1 codon. The β coefficients for each plot are in comparison to a control. For most of the amino acid and codon plots the control is Ala or an Alanine codon (GCA), respectively. For the EJC-enhanced **-2** position (E-F), the control is Arg or an Arg codon (AGA) for the amino acid and codon plots, respectively. For the amino acid plots, all β coefficients with a p-value < 0.01 are plotted in red and the red asterisk represents a stop codon. In the codon plots, all β coefficients with a p-value < 0.01 have a white asterisk, and “Stp” refers to stop codons.

Remarkably, for the linear model examining the effect of the amino acid at the -1 position on NMD activity, we found that Gly-PTC (the same context enriched in the gnomAD database; **Fig. 1**) had the strongest effect on NMD activity, increasing it by 41% compared to Ala (**Fig. 4C**). This increase was stronger than having a PTC at the -1 position. In contrast, Tyr-PTC had the least efficient NMD, weakening NMD by ∼25% compared to Ala-PTC (**Fig. 4C**). Thus, the effect range for amino acid identity at the -1 position was wider than that of the -2 position. Once again, all individual codons at the -1 position, except for one Leu codon, were grouped by the encoded amino acid (**Fig. 4D**; all codons compared to Ala-GCA).

For the EJC-enhanced reporters, there were fewer significant changes at both -1 and -2 positions (**Fig. 4E-H**). Note that the control feature for the -2 position is Arg or an Arg codon (**Fig. 4E-F**) and for the -1 position it is Ala or an Ala codon (**Fig. 4G-H**). Control features were picked based on amino acids or codons with β coefficients in the middle range of the model. At the -2 position, the NMD effect range for amino acids and codons was much smaller for EJC-enhanced NMD compared to EJC-independent NMD (**Fig. 4E-F**). Lys and Pro decreased NMD activity by ∼17% or increased it by ∼6%, respectively (**Fig. 4E**). When examining individual codons, only one Ala and one Lys codon had nominal but significant effects on NMD activity (**Fig. 4F**). At the -1 position, none of the amino acids had a significant impact on NMD (**Fig. 4G**). This trend was also present for codons, except for one Arg codon (AGG), which significantly weakened NMD by ∼20% compared to the Ala codon GCA (**Fig. 4H**). We also constructed linear models to examine the effect of the +4 nucleotide on NMD and found that it had some significant but nominal effects (**Supp. Fig. 2D**).

Taken together, the data show that the codons and amino acids at the -1 and -2 positions before the PTC influence NMD in a manner dependent on the mode of NMD (i.e. EJC-independent or EJC-enhanced). It is possible that the highly efficient EJC-enhanced NMD is insensitive to other modifying factors or that a narrow range of biological effects may not emerge as statistically significant in a large pool of ∼65k sequences.

To further explore the basis for the difference in the behavior of EJC-independent versus -enhanced modes of NMD, we switched to an independent, luciferase-based reporter system to test a limited number of sequence contexts and their relative impact on EJC-independent and EJC-enhanced NMD activity (Kolakada et al., 2024). In this reporter system, the control is Renilla luciferase and the NMD readout is provided by Firefly luciferase (**Supp. Fig. 3A**). As the -1 position of the PTC had the most significant impact on EJC-independent NMD activity in our MPRA, we checked its effect on EJC-independent and EJC-enhanced NMD activity in the luciferase reporter system. We varied the amino acid in the -1 position of either the PTC or the NTC, choosing Gly and Tyr, which were the contexts with the most efficient and least efficient NMD, respectively (**Fig 4C; Supp. Fig. 3A**).

We integrated these reporters into HEK293T cells and measured their steady state RNA levels with either DMSO or SMG1i treatment. RNA levels of NMD-reporters showed that Gly-NTC levels were lower than Tyr-NTC levels (**Supp. Fig. 3B**). However, this difference was not resolved by inhibiting NMD with SMG1i, suggesting that it was not due to NMD. Consistent with the results from the MPRA, Gly-PTC RNA levels were significantly lower than Tyr-PTC for the EJC-independent reporters. Inhibiting NMD resolved the difference, indicating that NMD was responsible for the discrepancy. Finally, there was no statistically significant difference between Gly-PTC and Tyr-PTC EJC-enhanced contexts, suggesting that EJC-enhanced NMD was indeed insensitive to sequence dependent modifying factors.

Measurements of RNA half-lives with Actinomycin D treatment of these reporters were consistent with the steady state results where only Gly-PTC and Tyr-PTC EJC-independent contexts had different half-lives (**Supp. Fig. 3C**). As the linearity of the curve ended after the first hour, we used the first 3 points to calculate half-lives across 3 replicates. Gly-PTC had a half-life of 0.53 ± 0.07 hours while Tyr-PTC had a half-life of 0.82 ± 0.13 hours, showing that the latter context was indeed more stable. Taken together, these data show that variation of the -1 amino acid before the PTC primarily influences EJC-independent NMD.

### Termination kinetics is a key determinant of NMD efficiency

To discover sequence-intrinsic factors that influence NMD, we constructed a random forest classifier trained to discriminate between constructs with high versus low NMD activity (**Fig. 5A**). We selected the 3000 sequences with the highest and lowest NMD activity for the EJC-independent and EJC-enhanced reporters. We then added sequence-intrinsic variables that could influence the classification of high and low NMD activity, including the identities of the codons as well as their properties such as usage frequency (GenScript), tRNA binding strength, and optimality (Bazzini et al., 2016; Grosjean & Westhof, 2016); the identities of the amino acids as well as their physical properties such as charge, solubility, dissociation constants, and hydropathy (ThermoFisher Scientific); and the GC and pyrimidine content of each of the sequences. As NMD is dependent on translation termination, which can vary in efficiency (Hellen, 2018; Lawson et al., 2021), we also included the rate of peptide release, which is a measure of the time taken for a release factor to trigger peptidyl-tRNA hydrolysis within the P-site (i.e. the -1 position of a stop codon) and the release of the polypeptide itself. These release rates vary based on the identity of the amino acid present within the P-site during termination. Since P-site amino acid dependent release rates are not available for eukaryotic systems, we used the prokaryotic peptide release rates (Pierson et al., 2016). Prokaryotes have two release factors RF1 and RF2 that each recognize a subset of the three stop codons, and therefore, we included the rates calculated for both RF1 and RF2 in our classifier.

**Figure 5.**
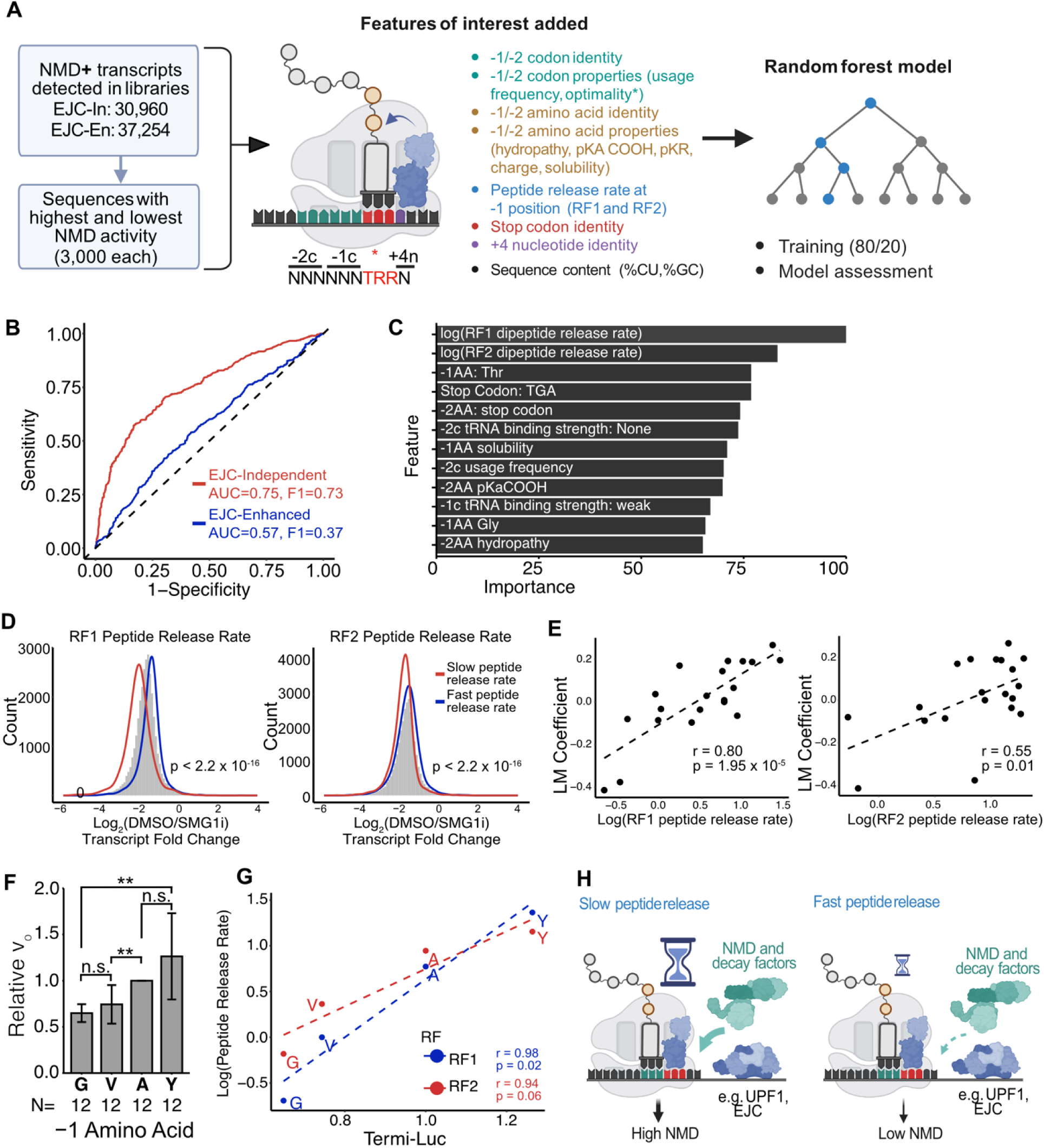
Peptide release rates predict NMD activity. **(A)** Data preparation and variables input into the random forest model. **(B)** ROC curves, AUC values, and F1 scores for the EJC-independent and EJC-enhanced classifiers. **(C)** Ranked feature importance for the EJC-independent model. **(D)** Density curves for sequences with slow and fast peptide release rates overlaid on the NMD activity distribution for the EJC-independent NMD+ sequences. Prokaryotic release rates used (Pierson et al., 2016), left: RF1 release rates, right: RF2 release rates. For RF1, slow release rates are defined as log (RF1 release rate) < –0.5, and fast release rates are defined as log (RF1 release rate) > 1. For RF2, slow release rates are defined as log (RF2 release rate) < 0, and fast release rates are defined as log (RF2 release rate) > 1.25. p, Wilcoxon test p-value **(E)** Scatterplot of the β coefficients for each -1 amino acid obtained for the EJC-independent reporters against the RF1 (left) and RF2 (right) peptide release rates. “r” refers to the Pearson’s correlation coefficient and “p” is the corresponding P-value. **(F)** Relative peptide release rates for each reporter as determined by Termi-Luc peptide release assay **(G)** A comparison of Termi-Luc determined eukaryotic release rates with Pierson et al prokaryotic release rates (Pierson et al., 2016). “r” refers to the Pearson’s correlation coefficient and “p” is the corresponding P-value. **(H)** A window of opportunity model for how peptide release rates influence NMD activity. A slower peptide release rate gives NMD factors a larger window to bind an act on a target, and vice versa.

We trained two separate classifiers via train (80%)/ test (20%) cross-validation for the EJC-independent and EJC-enhanced reporters using the same input variables. Classification performance for EJC-independent NMD activity was much better than for EJC-enhanced NMD activity (**Fig. 5B**), as indicated by the substantially higher area under the receiver operating characteristic curve and F1 scores for the EJC-independent (0.75 and 0.73, respectively) compared to the EJC-enhanced (0.57 and 0.37, respectively) classifiers. We concluded that only the classifier for EJC-independent NMD activity was robust and suitable for further interpretation (refer to **Supplementary Tables 1-2** for the confusion matrices and other model metrics). The poor performance of the EJC-enhanced classifier is consistent with EJC-enhanced NMD not being as influenced by sequence-dependent modifiers (**Supp. Fig. 3).** Of all the features that contributed to the EJC-independent model, prokaryotic release factor 1 (RF1) and release factor 2 (RF2) peptide release rates emerged as the most important predictor of NMD activity (**Fig 5C**).

Although these rates are measured from a prokaryotic context, the similarity in peptide release mechanisms between prokaryotes and eukaryotes suggests that the relative magnitudes of release rates may also apply to eukaryotic systems (Hellen, 2018). To visualize the effect of peptide release rates on NMD activity, we compared the subpopulation of sequences with the fastest and slowest dipeptide release rates versus all NMD targets (**Fig. 5D**). The two populations were well-separated for RF1 peptide release rates and less robustly for the RF2 peptide release rates (**Fig. 5D**). To examine this relationship more quantitatively, we compared the β coefficients of the effect of the peptide release rates for the -1 amino acid on NMD activity (**Fig 5E**). We observed a strong correlation for the RF1 peptide release rates (r = 0.8; p = 1.95 x 10^-5^) and a weaker but significant correlation for the RF2 peptide release rates (r = 0.55; p = 0.01) (**Fig 5E**). Taken together, these data suggest that peptide release rate may be a major factor contributing to the -1 amino acid-dependent impact on NMD activity.

The peptide release rate values used in the random forest classifier (**Fig. 5A**) came from a prokaryotic system and we wanted to determine whether they reflect the relative peptide release rate in a eukaryotic system (Pierson et al., 2016). To determine if the rate of peptide release varied based on the identity of the -1 amino acid in a eukaryotic system, we used the *in vitro* Termi-Luc assay (**Supp. Fig. 4A**). In this assay, we provide an excess of mutated eukaryotic release factor 1 (eRF1(AGQ)), which can recognize stop codons but are unable to induce peptidyl-tRNA hydrolysis (Frolova et al., 1999). eRF1(AGQ) binding thereby traps translation at the termination stage, allowing us to purify pre-termination ribosomal complexes (preTCs) from cells. PreTCs are then exposed to purified, functional release factors (working concentrations determined in **Supp. Fig. 4B**), and the real-time release kinetics of the synthesized NLuc are then measured to determine the translation termination rate of the transcript. We used this system to compare termination rates when Gly, Val, Ala, or Tyr codons (representing the full range of observed NMD efficiencies from high to low) were present at the - 1 position to the stop codon.

To check if the substitution of an amino acid residue affected NLuc activity, translation efficiency, or preTC stability of NLuc, control experiments were first performed. To determine whether NLuc activity was impacted by altering the last amino acid, we purified all tested peptide variants as recombinant proteins and determined their maximum luminescence. All NLuc peptide variants had the same activity (**Supp. Fig. 4C**). To determine whether translation efficiency was impacted by changing the last amino acid, we calculated the maximal derivative (slope) of the linear portion of the luminescence production curve. The identity of the last amino acid affected translation efficiency with Tyr having the lowest and Ala having the highest efficiency (**Supp. Fig. 4D**). To determine preTC stability, the rate of peptide release of each peptide variant was calculated without adding any release factors. These data showed that Gly had the most unstable preTC, while Tyr had the most stable preTC (**Supp. Fig. 4E**). Finally, we performed the Termi-Luc assay on each peptide variant starting with equal amounts of preTC complexes to account for the variability in translation efficiency and preTC stability. We found that Gly had the slowest release rate, and Tyr had the fastest release rate. This difference was statistically significant and correlated with high and low NMD activity, respectively (**Fig. 5F**). Finally, the release rates observed via the Termi-Luc assay were highly correlated with the values obtained in the prokaryotic system by Pierson *et al*. (**Fig. 5G**; r_RF1_ = 0.98; r_RF2_ = 0.94) (Pierson et al., 2016). These data show that peptide release rates indeed vary based on the identity of the last amino acid in a eukaryotic system, with glycine showing the slowest termination rate and tyrosine the fastest, among the amino acids tested. Taken together, our data supports a model where a slower peptide release rate offers a longer kinetic window for NMD factors to assemble and degrade a transcript, thus causing highly efficient NMD (**Fig. 5H**).

## DISCUSSION

NMD is often considered a binary outcome that relies on the presence of triggering factors downstream of the PTC. While recent results increasingly suggest that NMD is tunable (Jagannathan & Bradley, 2016; Lindeboom et al., 2019), the factors influencing this variability are incompletely understood. Our data support a window of opportunity model where the time of residence of the terminating ribosome determines the effectiveness of NMD on a given transcript (**Fig. 5H**). This model suggests that the ribosome serves as a temporal platform for the stepwise assembly of the NMD machinery, with slower peptide release rates enhancing NMD activity by extending this assembly window. While the precise temporal thresholds required for efficient NMD machinery recruitment remain to be determined, our findings suggest that termination kinetics, together with the RBP cues that trigger NMD, act as key modifiers of NMD activity.

The observation that led us to the window of opportunity model was the strong enrichment of glycine in the position immediately before the PTC in prematurely terminating transcripts. While Gly codons are the most enriched before a PTC, they are the least enriched before an NTC. This pattern is particularly striking among variants that are tolerant to loss of function in healthy individuals, where PTCs with Gly-PTC contexts show significant enrichment, while being notably absent among disease-causing variants. These enrichment patterns became logically coherent once we showed that Gly-PTC induces the most efficient EJC-independent NMD among all possible amino acid contexts at the -1 position and does so through its slow peptide release rate from the terminating ribosome. Interestingly, Gly is also a C-terminal degron, meaning that Gly-end proteins, truncated or otherwise, are more efficiently cleared (Koren et al., 2018; Lin et al., 2018). Thus, the preponderance of glycine codons at the ends of truncated proteins may have occurred through the course of evolution alongside processes that efficiently eliminate Gly-end proteins at both the RNA and protein levels.

The consistency of our model extends to other amino acids beyond glycine. Thr and Pro, which are also enriched before PTCs in healthy individuals and depleted before NTCs, support efficient NMD and exhibit slow peptide release rates. Conversely, Tyr, which shows the lowest NMD activity and fastest peptide release rate, is enriched before NTCs but not PTCs. PTCs with Tyr at the -1 of a PTC are depleted among variants tolerant to loss of function and enriched in disease contexts.

Apart from the -1 amino acid position, our massively parallel assessment of NMD activity allowed us to uncover other local sequence preferences for efficient NMD. Surprisingly, we found that the stop codon TGA caused 20% more efficient EJC-independent NMD than TAG. TGA also emerged as an important predictor of EJC-independent NMD in the random forest classifier. This finding appears to conflict with TGA’s known propensity for drug-induced readthrough, which typically stabilizes NMD targets (Tate & Mannering, 1996). This apparent paradox highlights the complex relationship between readthrough potential and intrinsic NMD activity. Recent work has shown that different drugs can induce varying degrees of readthrough depending on sequence context (Toledano et al., 2024), suggesting that the mechanisms governing natural termination efficiency and drug-induced readthrough may be distinct. Future studies combining structural biology approaches with biochemical assays could help elucidate how specific sequence elements influence both processes.

Our study highlights the power of human genetics to discover novel molecular mechanisms that can explain the real-world variability observed in the expression of genes containing nonsense variants. Our results point to a key role for the kinetics of translation termination in determining NMD activity, ultimately determining the efficiency with which prematurely terminating transcripts are cleared from the cell. The identification of sequence contexts that modulate NMD efficiency could inform the development of personalized therapeutic approaches for PTC-associated diseases. For instance, drugs targeting translation termination kinetics might be more effective in certain sequence contexts, suggesting the potential for context-specific therapeutic strategies. However, the success of such approaches will require a deeper understanding of how tissue-specific factors and varying cellular conditions influence the relationship between sequence context and NMD efficiency. The impact of amino acid identity upstream of the PTC on NMD activity and patterns of amino acid-PTC enrichment in healthy and diseased populations suggest that NMD has played a key role in shaping the evolution of nonsense variants. These findings can be used to better model the phenotypic outcomes of PTCs, and thus develop more effective strategies to counteract PTC-associated diseases.

## ACKNOWLEDGEMENTS

We thank Matthew Taliaferro for the gift of HEK293T cells with the loxP cassette. We thank Timothy Stasevich, Tatsuya Morisaki, and Laura White for exploratory studies on glycine’s impact on NMD. We thank Monkol Lek and Sander Pajusalu for helpful discussions on gnomAD analysis. We thank Srinivas Ramachandran, Olivia Rissland, and all members of the Jagannathan laboratory for insightful manuscript feedback. This work was supported by the RNA Bioscience Initiative (R. F.), AHA Award #831183 (D.K.), 5T32GM136444-03 (M.L.), Polish National Agency for Academic Exchange Bekker Program PPN/BEK/2019/1/00173 (M.P.S.), R35 GM119550 (J.H.), R35GM147025 (N.M.), TOPMed NHLBI fellowship (Z.C.A), Simons Foundation pilot award AGT011737 (Z.C.A and S.J.) University of Colorado School of Medicine Translational Research Scholars Program (S.J.), and the National Institutes of Health grant R35GM133433 (S.J.). The study of the effect of the stop codon 5’ context on the efficiency of translation termination was supported by the Russian Science Foundation grant 22-14-00279 (E.A.).

## AUTHOR CONTRIBUTIONS

Conceptualization (D.K., R.F., S.J.); Data curation (D.K., R.F., A.E.C., S.J.); Formal analysis (D.K., R.F., Z. C. A., S.J.); Funding acquisition (D.K., E.A., Z. C. A., S.J.); Investigation (D.K., R.F., N.B., A.S., M.L., A.E.C., S.J.); Project administration (S.J.); Software (D.K., R.F., M.P.S., M.A.C., N.M., S.J.); Supervision (J.H., N.M., E.A., S.J.); Validation (D.K., A.E.C.); Visualization (D.K., R.F., A.E.C., Z. C. A., S.J.); Writing – original draft (D.K., S.J.); Writing – review and editing (D.K., R.F., A.E.C., M.A.C, M.P.S., J.H., N.M., E.A., S.J.).

## METHODS

### RESOURCE AVAILABILITY

#### Lead contact

Further information and requests for resources and reagents should be directed to and will be fulfilled by the lead contact, Sujatha Jagannathan.

#### Materials availability

All unique reagents generated in this study are available from the lead contact with a completed Material Transfer Agreement.

#### Data and code availability

- All sequencing data have been deposited at GEO (GSE261151) and will be publicly available as of publication.
- All original code is available at the GitHub repository: https://github.com/jagannathan-lab/2024-kolakada_et_al.
- Any additional information required to reanalyze the data reported in this paper is available from the lead contact upon request.

## METHOD DETAILS

### gnomAD and ClinVar analysis

Rare (frequency < 0.01) SNP variants from gnomAD v2.1.1 exome database (Karczewski et al., 2020) that passed gnomAD’s built-in random forest classification quality filtering were analyzed, filtering for “Stop-gained” consequence via bcftools (https://github.com/samtools/bcftools; (Danecek et al., 2021)). Using “cDNA_position” field pre-annotated by VEP (McLaren et al., 2016), the sequence and codons surrounding variants from Gencode (Frankish et al., 2023) v19 transcript FASTA file was then extracted using bedtools getfasta (Quinlan & Hall, 2010). For other contexts such as normal stop codons, custom code (found in: https://github.com/jagannathan-lab/2024-kolakada_et_al) was used to extract sequence and codon information using Biostrings (https://github.com/Bioconductor/Biostrings) and BSgenome (https://github.com/Bioconductor/BSgenome) in R. Codon enrichment is calculated as observed frequency of a particular codon/amino acid normalized against normal occurrence frequency of the codon/amino acid in the last 10 positions of all normal coding genes, similar to Koren et al (Koren et al., 2018). Gene-level loss of function scores were retrieved from gnomAD LoF constraint scores. Clinvar data (Landrum et al., 2014) from 2022/07/30 was processed similarly to gnomAD, but with sequence extraction done in R.

### TOPMed Allele-specific expression analysis

“Freeze 1.1RNA TOPMed cis-e/sQTL results were generated in a collaboration between the TOPMed Informatics Research Center, TOPMed Multi-Omics working group, and the TOPMed parent studies contributing RNA-seq and distributed to TOPMed investigators. For the accurate calculation of allele specific expression (ASE) from RNA-Seq data, we used STAR aligner’s WASP functionality (van de Geijn et al., 2015) to mitigate the reference bias while mapping the RNA-Seq reads to the reference genome. Then we used GATK ASEReadCounter version (v3) (https://gatk.broadinstitute.org/hc/en-us/articles/360037428291-ASEReadCounter) (Castel et al., 2020) to extract allele specific expression for truncating variants. NMD efficiency was then calculated as reference read allele count/total allele count.

### Cell lines and culture conditions

HEK293T cells (female) were obtained from ATCC (CRL-3216; RRID:CVCL_0063). HEK293T Lox2272/LoxP cells were obtained from Taliaferro laboratory. All cell lines were determined to be free of mycoplasma by PCR screening. HEK293T and HEK293T cells with Lox2272/LoxP cells were maintained in Dulbecco’s Modified Eagle Medium (DMEM) (Thermo Fisher Scientific) supplemented with 10% EqualFETAL (Atlas Biologicals).

### Cloning

The NMD reporter plasmids have been previously described (Kolakada et al., 2024). For individual luminescent reporters containing Gly, and Tyr amino acids before the PTC, oligos containing these sequences (Integrated DNA Technologies) and matching the EcoRI (NEB,

R3101S) and XhoI (NEB, R0146S) restriction sites were synthesized. These oligos were annealed and ligated using the Quick Ligation Kit (NEB, M2200L) into EJC-independent and EJC-enhanced fluorescent and luminescent backbones digested with EcoRI and XhoI. To make the NMD**–** reporters for each sequence, each EJC-independent reporter was digested with EcoRI and MfeI (NEB, R3589S), the sticky ends were filled in using Klenow Fragment (NEB, M0212S) and the blunt ends were ligated together using the Quick Ligation Kit (NEB), removing the GFP 3′ UTR. The oligos synthesized for each sequence tested are as follows.

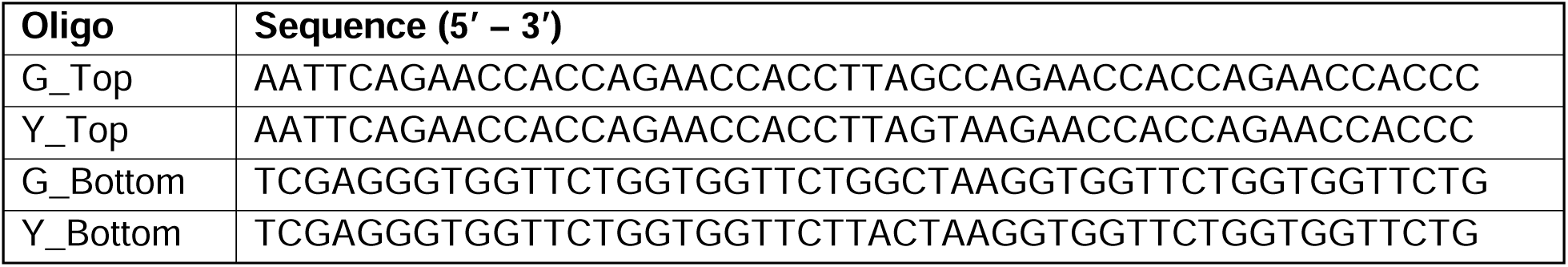

### MPRA plasmid library

An oligo library (Eurofins) of sequence CTA GCA AAC TGG GGC ACA GCC TCG AGG GTG GTT CTG GTG GTN NNN NNT RRN GTG GTT CTG GTG GTT CTG AAT TCG ACT ACA AGG ACC ACG ACG G was synthesized, where N refers to equal proportions of the A, C, G, and T nucleotides, and R refers to equal portions of the A and G nucleotides. This library was resuspended in water to 100 μM. The oligo libraries were then filled in (1 amplification cycle) using the Q5 High-Fidelity DNA Polymerase (NEB, M0492S) according to manufacturer’s instructions, using 100 μmols of oligos, and 100 μmols of reverse primer (sequence: CACCGTCGTGGTCCTTGTAGTC), in 2× PCR reactions of 25 μL each. After amplification, the PCR reaction was digested with Exonuclease I, at 37°C for 3 h to digest any remaining single stranded DNA. The DNA was purified using AMPure XP beads (Beckman, A63882), as per manufacturer’s instructions.

The EJC-independent and EJC-enhanced fluorescent reporter backbones were linearized using EcoRI (NEB) and XhoI (NEB) at room temperature overnight. Digested plasmid DNA was gel purified using the NucleoSpin Gel and PCR Clean-up kit (Takara Bio, 740609.250). The oligo library was then cloned into the digested backbones using the Gibson Assembly Master Mix (NEB, E2611L) using an insert:vector molar ratio of 7:1 for the EJC-independent backbone and 3:1 for the EJC-enhanced backbone. The reactions were incubated at 50°C for 1 h. The reactions were then purified using AMPure XP beads to remove excess salts and a total of 200 ng of plasmid DNA was transformed into MegaX DH10B T1R Electrocompetent Cells (ThermoFisher, C640003), using a Biorad GenePulser electroporator. The transformed cells were recovered in the recovery medium provided with the electrocompetent cells at 37°C for 1 h, then plated on 33 pre-warmed 15 cm Luria broth (LB) agar-Carbenicillin (RPI, C46000-5.0) plates and incubated at 37°C overnight. The next day colonies were collected by scraping plates, spun-down, and then midi-prepped (NucleoBond Xtra Midi EF kit, 740420.50) to extract the plasmid DNA.

### Generating stable cell lines

To create the EJC-independent and EJC-enhanced reporter cell lines HEK293T cells containing a Lox2272 and LoxP enclosing a blasticidin cassette (courtesy Taliaferro Lab, CU Anschutz), were co-transfected with 97.5% NMD reporter plasmid library and 2.5% Cre-plasmid (Addgene plasmid #27493). These transfections were performed in 60-70% confluent 10 cm plates using 11,800 ng of plasmid library and 295 ng of Cre plasmid per plate for 20 plates. Twenty-four hours after transfection, each plate of cells was split equally into two new plates and puromycin (2 μg/mL) was added to the media to begin selection. Puromycin selection lasted between 1-2 weeks, during which the media was replaced with fresh media supplemented with puromycin every 1-2 days and cells were consolidated into fewer plates when their confluency got too low. Once selection was complete, cells were frozen down in medium containing 10% DMSO. These cells were thawed at a later time for the MPRA.

The same method was used to create the luciferase reporter cell lines, however transfections were performed in 50% confluent 6-well plates using 2437.5 ng of each plasmid and 62.5 ng of Cre plasmid per well.

### Targeted sequencing of MPRA libraries

To obtain MPRA sequencing libraries, frozen EJC-independent and EJC-enhanced reporter cell lines were thawed. These cells were split into 6, 15 cm plates, 3 of which were treated with DMSO and the other 3 treated with 0.5 μM SMG1i (technical replicates). These plates were harvested for RNA in TRIzol Reagent (ThermoFisher Scientific, 15596018) after 24 hrs. TRIzol extractions were performed for RNA as per manufacturer’s instructions. A DNase digest was conducted on 10 μg of RNA from each sample using the Turbo DNA-free kit (ThermoFisher Scientific, AM1907), including reaction inactivation. Following this, 2 μg of digested RNA was used to make cDNA using Superscript II Reverse Transcriptase in duplicate 20 μL reactions (ThermoFisher Scientific, 18064014).

For library preparation, each cDNA sample was split into 20 PCR reactions (2 μL cDNA/PCR) and amplified using reporter-specific forward and reverse primers with UMIs (forward primer: ACACTCTTTCCCTACACGACGCTCTTCCGATCTGCAGACTTCCTCTGCCCTC; reverse primer: AGACGTGTGCTCTTCCGATCTNNNNNNNNtggggcacagcctcga, where the Ns refer to the nucleotides for the UMIs). PCR reactions were performed using Kapa HotStart PCR Kit (Roche, KAPA KK2502) using an annealing temperature of 69°C and 32× cycles. The 20 PCR reactions were pooled and 100 μL of all reactions were purified using AMPure XP beads. A second PCR reaction was performed on 25 ng of the purified product with primers containing Illumina adaptors. Once again, the Kapa HotStart PCR Kit was used, this time with an annealing temperature of 65°C and 8x cycles. Reactions were purified using AMPure XP beads and samples were run on an agarose gel to verify the library size. Libraries were sequenced using the Illumina sequencing platform NovaSeq 6000 (paired-end 2×150 cycles). A total of 50 million and 30 million reads were requested for the EJC-independent and EJC-enhanced libraries, respectively.

### MPRA data analysis

To process the MPRA sequencing libraries, the UMI-tools repository (https://github.com/CGATOxford/UMI-tools) was used to add the UMIs to the name of each read in the FASTQ files (Smith et al., 2017). Following this, Cutadapt was used to trim the 5′ and 3′ends of the reads in a sequence dependent manner, up to the 10 nt context of interest (Martin, 2011); ∼25% of reads were lost in this process (https://github.com/marcelm/cutadapt). SeqPrep was used to merge sequencing reads 1 and 2, only accepting reads that perfectly align (https://github.com/jstjohn/SeqPrep); ∼5% of reads were lost in this process. Bowtie was used to map the reads to a reference file of all sequence contexts (Langmead et al., 2009); 1% of reads were lost in this process. Samtools was used to create a BAM file of mapped reads, followed by deduplicating the BAM files based on UMIs using UMI-tools and converting the BAM files to BED files using samtools again (Li et al., 2009). Around 10% of reads were lost upon deduplication of UMIs. A custom python script was used to remove reads on the negative strand and to get a bed file of reads in the correct orientation.

The NMD activity of an individual reporter transcript was the log2-fold change in between DMSO-treated cells to SMG1i-treated cells calculated with DESeq2 (Love et al., 2014). The reporter transcripts that do not undergo NMD, i.e. Trp codon rather than a stop codon at the PTC position, were used similar to housekeeping normalization controls within DESeq2. Both linear models and random forest classifiers were performed in R. Random forest models were built utilizing the caret R package v. 6.0.92 (Kuhn, 2008). First, features with variance close to zero were eliminated, after which the linear correlation for all numeric features was calculated. For the features with correlation ≥ 0.85 one feature was randomly selected, and the others discarded from the model. Data were divided into training and validation set using 0.8:0.2 ratio. Hyperparameters tuning was performed for train set using 5x train-test cross validation for the following model parameters: mtry, maxnodes and ntree. Performance of final model was measured using F1 scores and on validation set. The code and packages used for these analyses can be found in this repository: https://github.com/jagannathan-lab/2024-kolakada_et_al.

### RNA extraction and RT-qPCR

For RT-qPCR used to determine steady state RNA levels of the luciferase reporters, RNA was extracted via the Qiagen RNeasy Plus Mini Kit (Qiagen 74136) as per manufacturer’s instructions. DNase digestion was performed with Turbo DNA-free Kit (Thermofisher Scientific, AM1907) as per manufacturer’s instructions on 5 μg of extracted RNA. This was followed by cDNA synthesis using the SuperScript III First-Strand Synthesis kit, using random hexamers. A no-RT sample was included as a control to make sure there was no genomic DNA contamination. The cDNA was then diluted 1:4 and 2 μL used per 10 μL qPCR reaction. qPCR was performed using PrimeTime Gene Expression Master Mix (Integrated DNA Technologies, 1055772). and a probe and primer mix specific to firefly luciferase and renilla luciferase to a final concentration of 0.25 μM and 0.5 μM for the probe and primers, respectively. These were the sequences of the primers and probes:

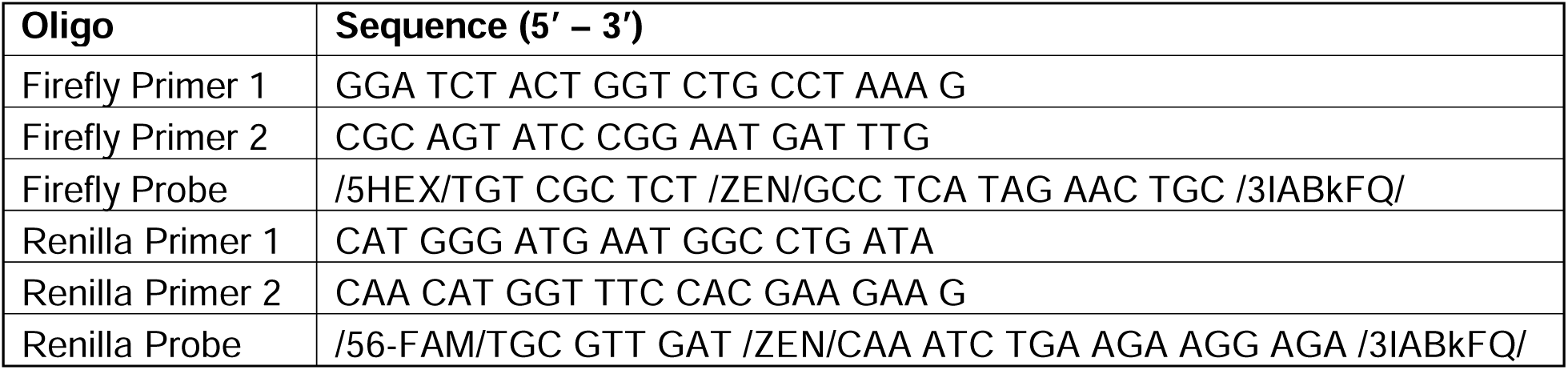

Reactions were set up in triplicate per sample and plated in 384-well plates. The plates were then run on the OPUS Bio-Rad qPCR machine (Bio-Rad, 12011319) using the 2-step Amplification and melting curve protocol as per PrimeTime manufacturer’s instructions. For the RNA level analysis, the Livak method (Livak & Schmittgen, 2001) was used normalizing Firefly to Renilla. Mean RNA levels of 3 different transfections or 3 technical replicates of transductions were plotted, respectively. Error was calculated using standard deviation.

### Half-life measurements using transcription inhibition

Half-life was determined using transcription inhibition followed by RT-qPCR. HEK293T Lox2272/LoxP cell lines stably expressing luciferase NMD- or NMD+ reporters were treated with 5 ug/mL actinomycin D (Sigma-Aldrich A9415). RNA was isolated at 0, 0.5, 1, 2, 4, and 8 hour time points using the RNeasy Plus Mini Kit (Qiagen 74136) and used as a template for DNase treatment and cDNA synthesis. Each data point had 3 technical replicates. Reporter mRNA levels were then quantified by qPCR as described in the qPCR section. To calculate the half-lives, linear models were constructed of each replicate using the *lm* function in R using the formula (formula = log (Fold-change) ∼ Time). The L coefficient measured degradation rate, which was used in the equation ln(2)/L to calculate the half-lives. Only the first 3 time points were used in the half-life calculations since linearity of the curve ended there. Error was calculated using the standard deviation across replicates.

### Luminescent reporters for *in vitro* assays

A pNL-globine vector derived from pNL1.1 vector (Promega), with a β-globin 5′UTR addition before nanoluciferase (NLuc) coding sequence was used (Shuvalov et al., 2021). Additional constructs were created based on pNL-globin, encoding NLuc with various substitutions of the codon before the stop codon using the QuikChange Site-Directed Mutagenesis Kit (Agilent Technologies, cat. 200518-5) was used. Primers were selected in the manufacturer’s recommended web-based QuikChange Primer Design Program (www.agilent.com/genomics/qcpd). For obtaining mRNA, the fragment of the plasmid were amplified using RV3L (CTAGCAAAATAGGCTGTCCCCAG) and FLA50 (TTTTTTTTTTTTTTTTTTTTTTTTTTTTTTTTTTTTTTTTTTTTTTTTTTAACTTGTTTATTGCAGCTTATAATGG) primers, as described in Shuvaolv et al., 2021 (Shuvalov et al., 2021). Templates were run-off transcribed with the T7 RiboMAX ™ Large Scale RNA Production System (Promega, P1320) kit according to the manufacturer’s protocol. The mRNA was then purified sequentially by isolation in acidic phenol, precipitation with 3 M LiCl followed by 80% ethanol wash.

### Expression and purification of eRF1 and eRF3

Recombinant eRF1 was expressed from the plasmid pET-SUMO-eRF1. To obtain pET-SUMO-eRF1, the eRF1 coding sequence was amplified from the plasmid pET23b-eRF1 (Frolova et al., 2002) using primers petSUMO_eRF1_F (GAGAACAGATTGGTGGTATGGCGGACGACCCCAG) and petSUMO_eRF1_R (CCGAATAAATACCTAAGCTCTAGTAGTCATCAAGGTCAAAAAATTCATCGTCTCCTCC). eRF1 was expressed in E. coli BL21(DE3) cells. The lysate was prepared by ultrasonification of the pelleted cells in a buffer composed of 20 mM Tris-HCl pH 7.5, 500 mM KCl, 10% glycerol, 0.1% Triton X-100, 0.5 mM PMSF, and 1 mM DTT. Following this, His-SUMO tag was cleaved by His-tagged Ulp1 protease. Untagged eRF1 was further purified by anion-exchange chromatography (HiTrap Q HP, Cytiva). Fractions enriched with eRF1 were collected, dialyzed in storage buffer, frizzed in liquid nitrogen, and stored at −70 °C. Recombinant eRF3A cloned into baculovirus vector EMBacY from a MultiBac expression system was expressed in the insect cell line Sf21. Following this, recombinant proteins were purified using Ni-NTA agarose and ion-exchange chromatography, as described previously (Ivanov et al., 2016).

### Expression, purification, and determination of luminescence or recombinant nanoluciferase

To obtain recombinant NLuc with varying amino acids in the position -1 to the stop codon, corresponding constructs of pNL-globine were created using the petSUMO vector (Invitrogen, cat. K30001). Each vector was assembled via Gibson Assembly (NEB, cat. E2611L) using PCR products of the petSUMO backbone (forward primer: AGCTTAGGTATTTATTCGGCGCAAAGTG; reverse primer: ACCACCAATCTGTTCTCTGTGAGC) and pNL-globine inserts (forward primer: GAGAACAGATTGGTGGTATGGTCTTCACACTCGAAGATTTCGTTGG; reverse primer: CCGAATAAATACCTAAGCTTACGCCAGAATGCGTTCGCA) amplified using Q5 High-Fidelity DNA Polymerase (NEB, cat. M0491L). The resulting plasmids were expressed in *E. coli* BL21(DE3) and His-SUMO-Nluc was purified using Ni-affinity chromatography. Recombinant 6xHis-ULP1 protease was added to purify 6xHisSUMO-Nluc and incubated for 2 hours. The 6xHisSUMO fragment and 6xHis-ULP1 were then removed with Ni-NTA agarose to obtain purified Nluc without tags. 0.1 femtomole of recombinant Nluc was incubated in storage buffer, with the addition of BSA up to 500 µg/mL and 0.5 % NanoGlo (Promega). Luminescence was measured at 30°C using a Tecan Infinite 200 Pro (Tecan, Männedorf, Switzerland) in a 40 min period. The luminescence of NLuc was calculated as a maximum of relative luminescence units (RLU_max_).

### Purification of the preTC-NLuc and Termi-Luc Assay

For the Termi-Luc assay, preTCs translating NLuc (preTC-NLuc) were purified using previously published methods (Shuvalov et al., 2021; Susorov et al., 2020). 100% RRL lysate was preincubated in a mixture containing 1 mM CaCl_2_ and 3 U/µL Micrococcal nuclease (Fermentas) at 30°C for 10 min, followed by the addition of EGTA to a final concentration of 4 mM. The lysate was then diluted to 70% (v/v) and supplemented with 20 mM HEPES-KOH (pH 7.5), 80 mM KOAc, 0.5 mM Mg(OAc)_2_, 0.3 mM ATP, 0.2 GTP, 0.04 mM of each of 20 amino acids (Promega), 0.5 mM spermidine, 0.45 µM aminoacylated total rabbit tRNA, 10 mM creatine phosphate, 0.003 U/µL creatine kinase (Sigma), 2 mM DTT, and 0.2 U/µL Ribolock (ThermoFisher) (70% RRL mix).

For preTC-NLuc assembly, 220 µL of 70% RRL mix was preincubated in the presence of 1.7 μM ERF1 G183A mutant (Alkalaeva et al., 2006) at 30° C for 10 min, followed by the addition of 10.5 pmol of NLuc mRNA. The mixture was incubated at 30°C for 40 min. The KOAc concentration was then adjusted to 300 mM and the mixture was layered on 5 ml of a 10–35% linear sucrose gradient in a buffer containing 50 mM HEPES-KOH, pH 7.5, 7.5 mM Mg(OAc)_2_, 300 mM KOAc, 2 mM DTT. The gradient was centrifuged in a SW55-Ti (Beckman Coulter) rotor at 55 000 rpm (367 598 g_max_) for 1 h. The gradient was fractionated in 15 fractions of 150 μL from bottom to top and the remaining sucrose was collected separately. Fractions of the first peak, enriched with the preTC-Nluc were analyzed by the Termi-Luc assay, merged, flash-frozen in liquid nitrogen, and stored at −70°C.

A peptide release assay with the preTC-NLuc (Termi-Luc) was performed as previously described with some modifications (Susorov et al., 2020). Peptide release was conducted in a solution containing 1.5 pM preTC-Nluc, 45 mM HEPES-KOH pH 7.5, 1.4 mM Mg_2_OAc, 56 mM KOAc pH 7.0, 1 mM DTT, 177 µM spermidine, 1.5 % (w/w) sucrose, 0.8 mM MgCl_2_, 0.2 mM GTP supplemented with equimolar MgCl_2_, and 0.5 % NanoGlo (Promega), in the absence or the presence of release factors eRF1 and eRF3a at various concentrations. Luminescence was measured at 30°C using a Tecan Infinite 200 Pro (Tecan, Männedorf, Switzerland) for 12 min. The translation efficiency was calculated as the maximal derivative of the growing linear section of the luminescence curve (v_0_, RLU/min). For comparison of different preTCs, a single working concentration of 3 nM of eRF1-eRF3A was chosen in the linear section. The instability of the preTC-NLuc was evaluated in Termi-Luc assay in the absence of release factors.

The amount of preTC-Nluc was calculated as a maximum of relative luminescence (RLU_max_). To determine concentration of preTC-Nluc, 1.25 µL of preTC sample were incubated in the presence of excess of eRF1 and eRF3a (100 nM). Luminescence was measured at 30°C using a Tecan Infinite 200 Pro (Tecan, Männedorf, Switzerland) for 40 min. PreTC concentration was considered in the translation termination rate experiments.

### Translation of NLuc in RRL lysates

For translation of NLuc, 19 μL of 70% RRL lysate prepared as described in Termi-Luc section, was mixed with 0.19 μL of NanoGlo and 0.25 pmol of NLuc mRNA. Luminescence was measured at 30°C using a Tecan Infinite 200 Pro (Tecan, Männedorf, Switzerland) for 60 min. The translation efficiency was calculated as a maximal derivative of the growing linear section of the luminescence curve (v_0_, RLU/min).

## QUANTIFICATION AND STATISTICAL ANALYSIS

### Data analysis, statistical tests, and visualization

All experiments were performed with a sample size of at least n=3. For transient transfections, each replicate represents an independent transfection. For stably integrated cells, the same cell line was plated in 3 different wells, from which RNA and protein were harvested. Statistical significance between various populations was calculated using a student’s t-test in R and p-values were two-sided (*p<0.05, **p<0.005, ***p<0.0005). Statistical details of specific experiments can be found in the Results, Methods, and/or Figure Legends. Plots were generated using R plotting functions and/or the ggplot2 package. The exact code and packages used for these analyses can be found in this repository: https://github.com/jagannathan-lab/2024-kolakada_et_al.

## SUPPLEMENTAL INFORMATION

### SUPPLEMENTARY FIGURES

**Figure S1.**
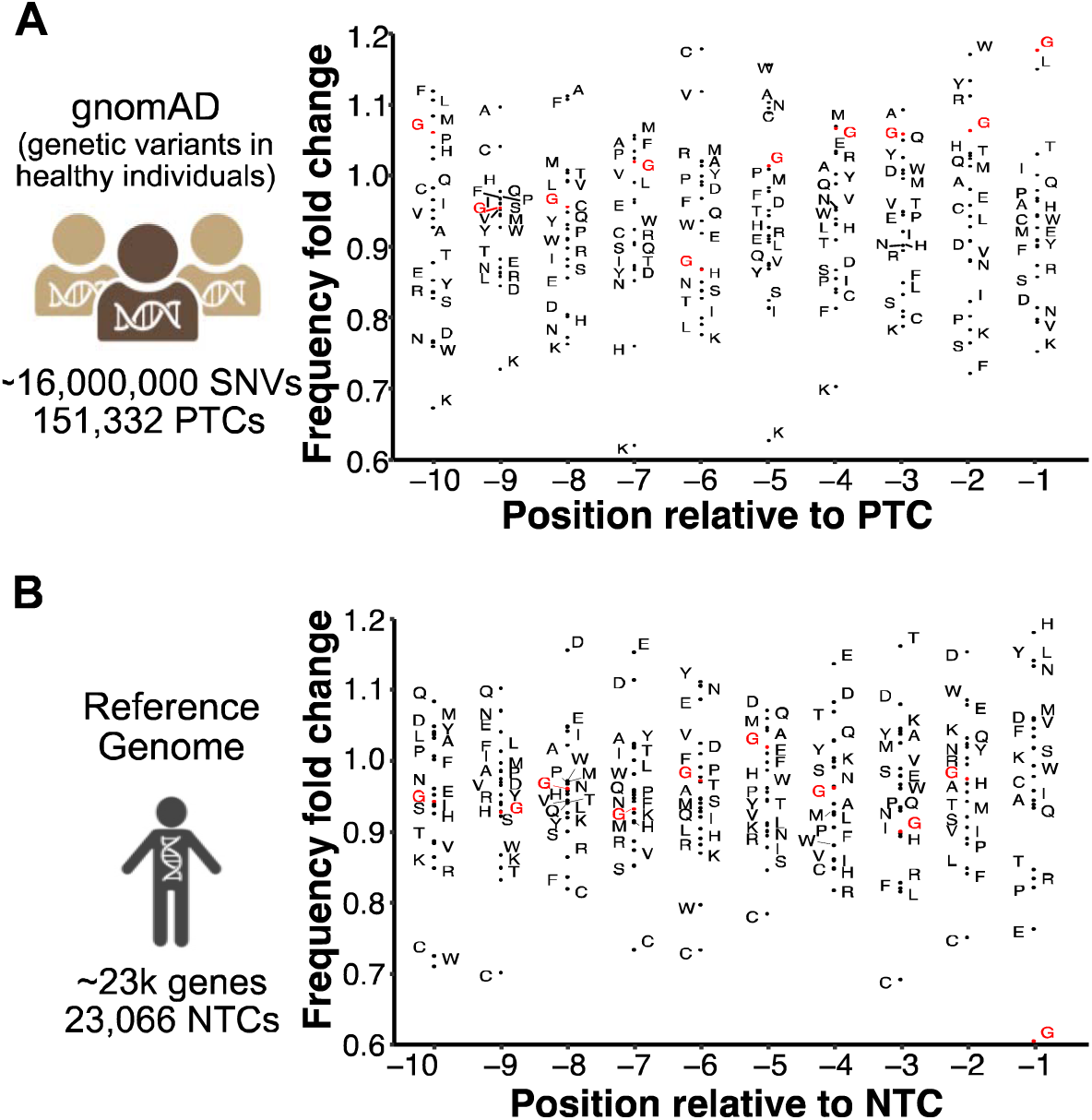
Amino acids enriched from the -10 to the -1 position before a termination codon. (**A**) Left: Schematic to indicate PTCs analyzed in the gnomAD database, right: amino acid enrichment before the PTC. (**B**) Left: Schematic to indicate NTCs analyzed in the reference genome, right: amino acid enrichment before the NTC. Gly is colored in red.

**Figure S2.**
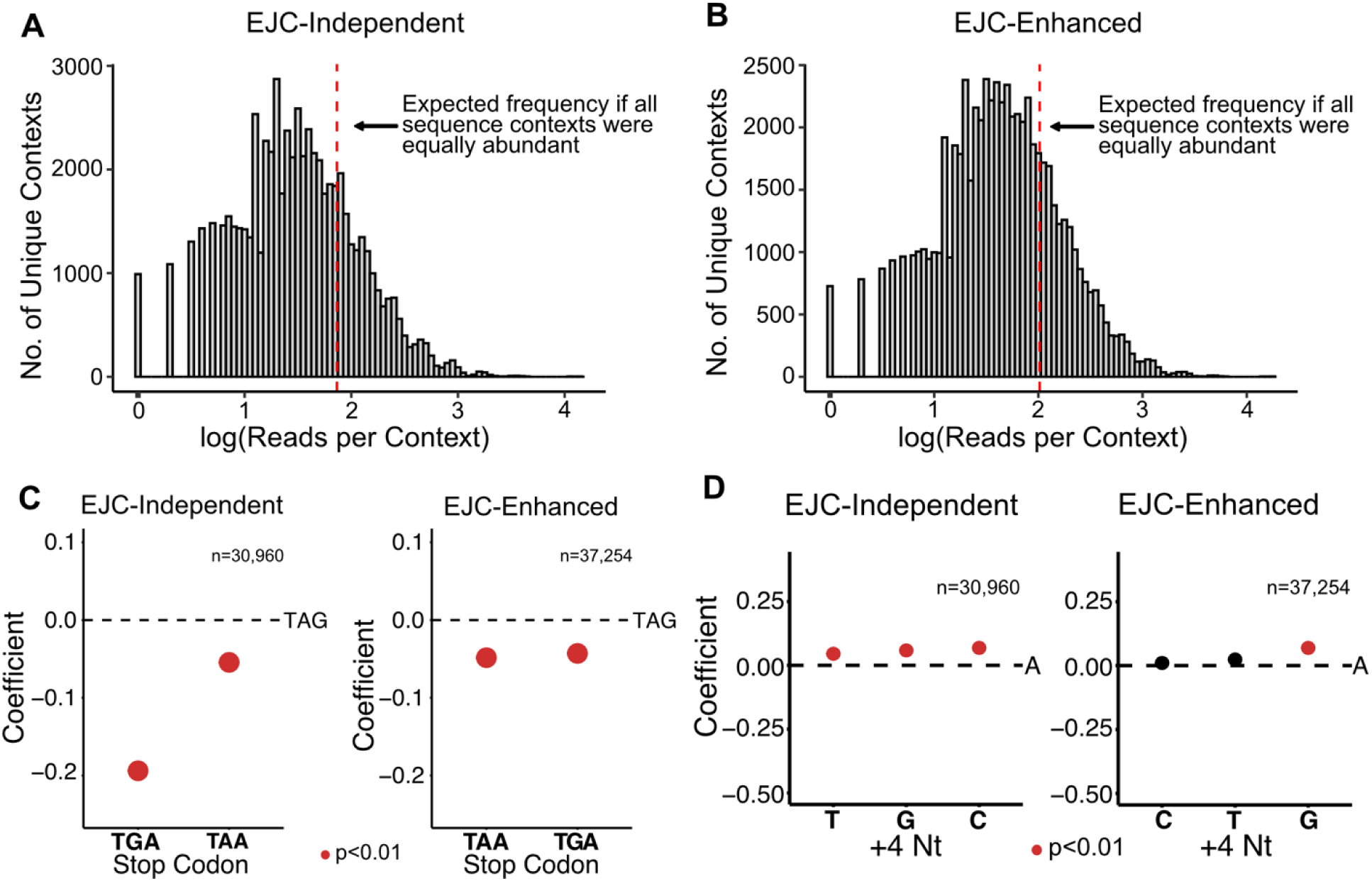
MPRA library representation. (**A**) Plasmid library representation for the EJC-independent NMD**+** reporter library. (**B**) The same as (A) but for the EJC-enhanced NMD**+** reporter library. (C) Effect of stop codon identity on EJC-independent and EJC-enhanced NMD without TGG. (D) Dot plots of the β coefficient representing the effect of the +4 nt on EJC-independent and EJC-enhanced NMD activity.

**Figure S3.**
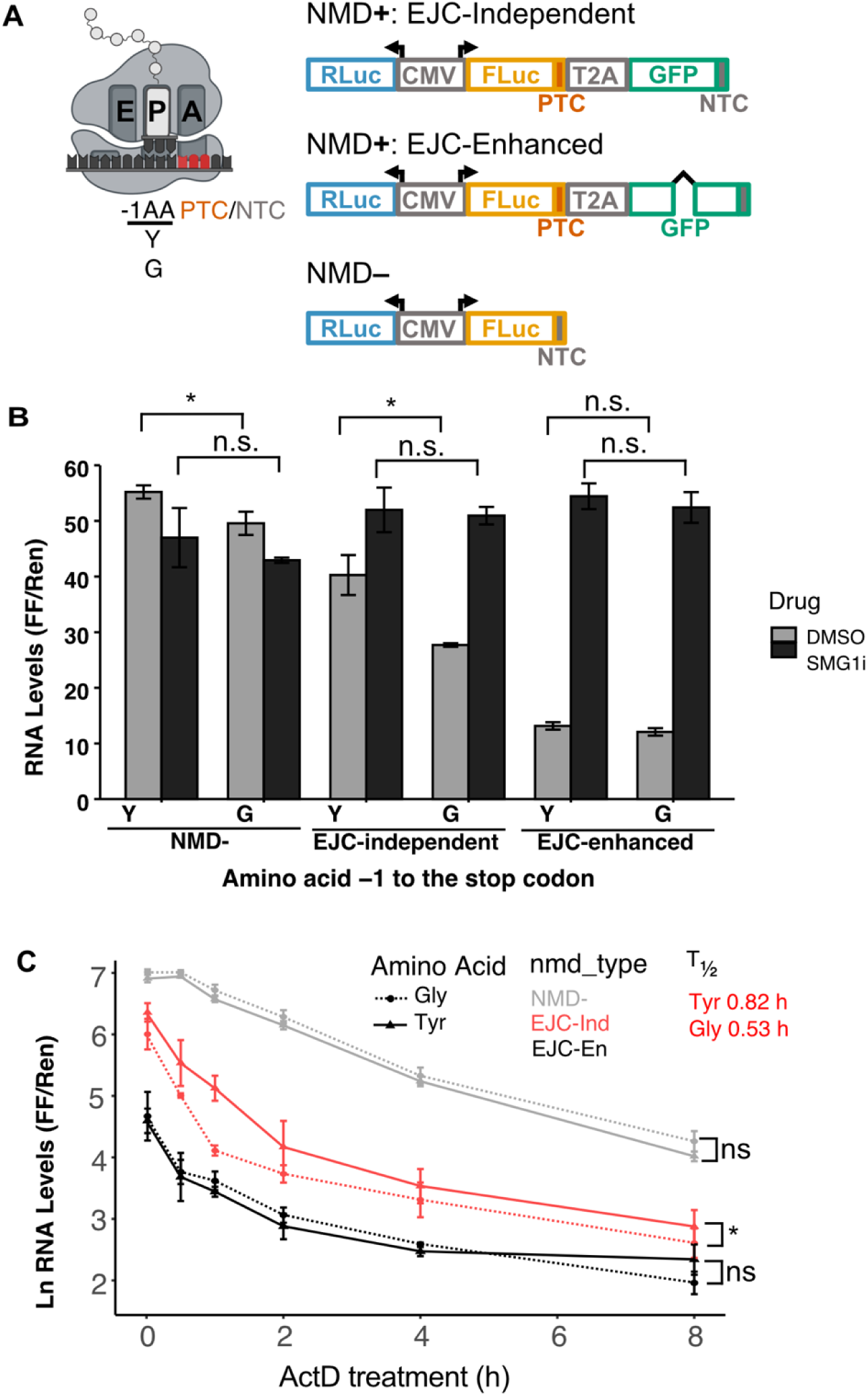
The codon identity at the -1 position of the PTC alters NMD activity of EJC-enhanced luminescence-based reporters. (A) Schematic representation of luminescence based NMD reporters used (right) as well as the amino acids tested in the -1 position to the PTC/NTC (left). (B) RNA level of luciferase reporters in EJC-independent, EJC-enhanced, and NMD– contexts. (C) RNA half-life estimation after transcription arrest with Actinomycin D treatment. Normalized reporter RNA levels at 0, 0.5, 1, 2, 4, and 8 hours post-actinomycin D treatment estimated across three replicates. Half-life calculated for the first three time points, when linearity is maintained.

**Figure S4.**
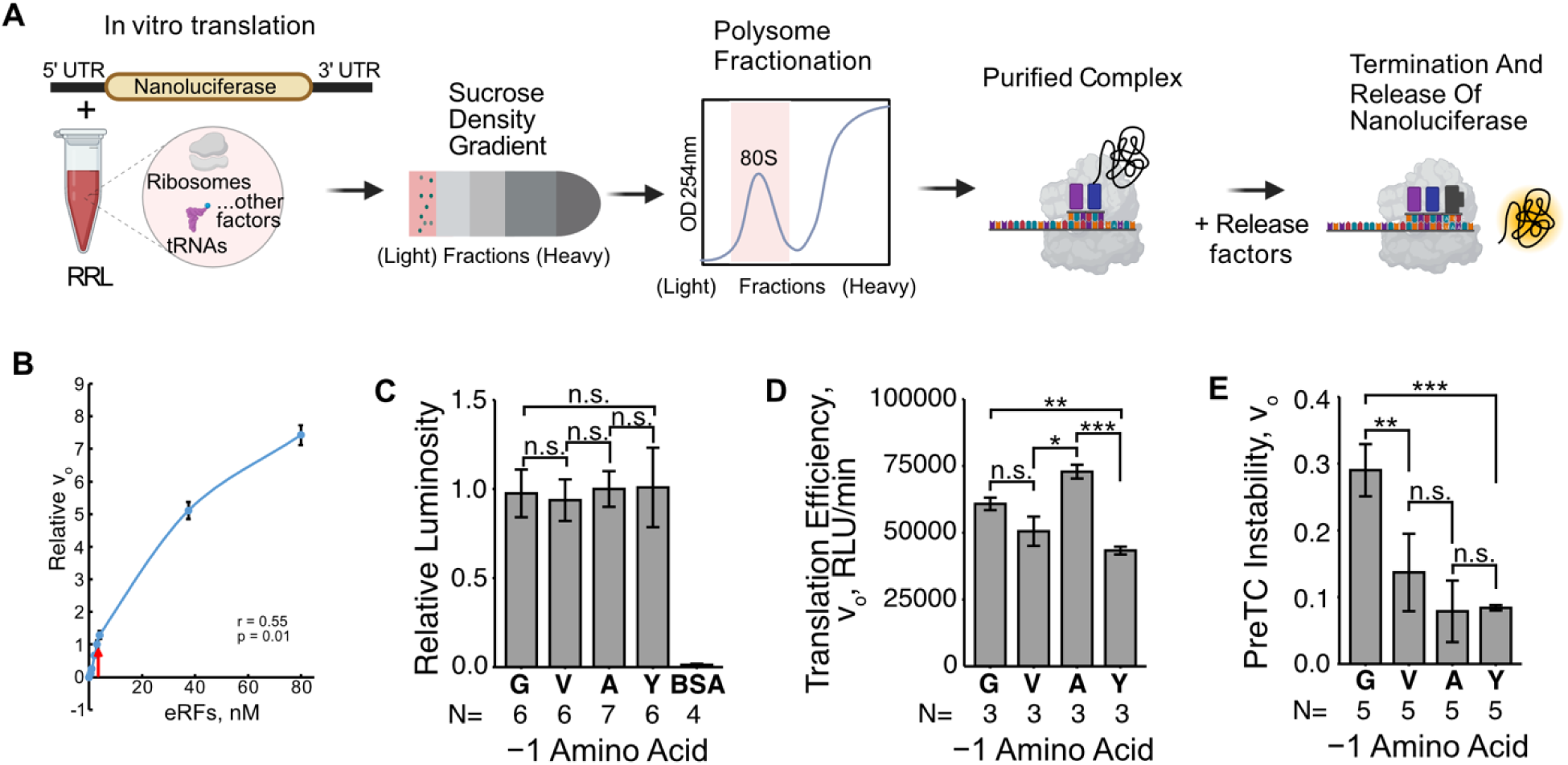
Peptide release rate estimation in an in vitro eukaryotic system via the Termi-Luc assay. **(A)** Schematic representation of the Termi-Luc peptide release assay. This assay was performed for nanoluciferase reporters with different amino acids -1 to the stop codon. **(B)** Determining the optimum concentration of release factors to use for the Termi-Luc assay. **(C)** Tested *in vitro* luminescence activity of recombinant NLuc variants. **(D)** Translation efficiency of each reporter in the *in vitro* system. **(E)** Relative rate of peptide release when no release factors are added to the termination reaction i.e., preTC instability.

#### SUPPLEMENTARY TABLES

**Table 1:**
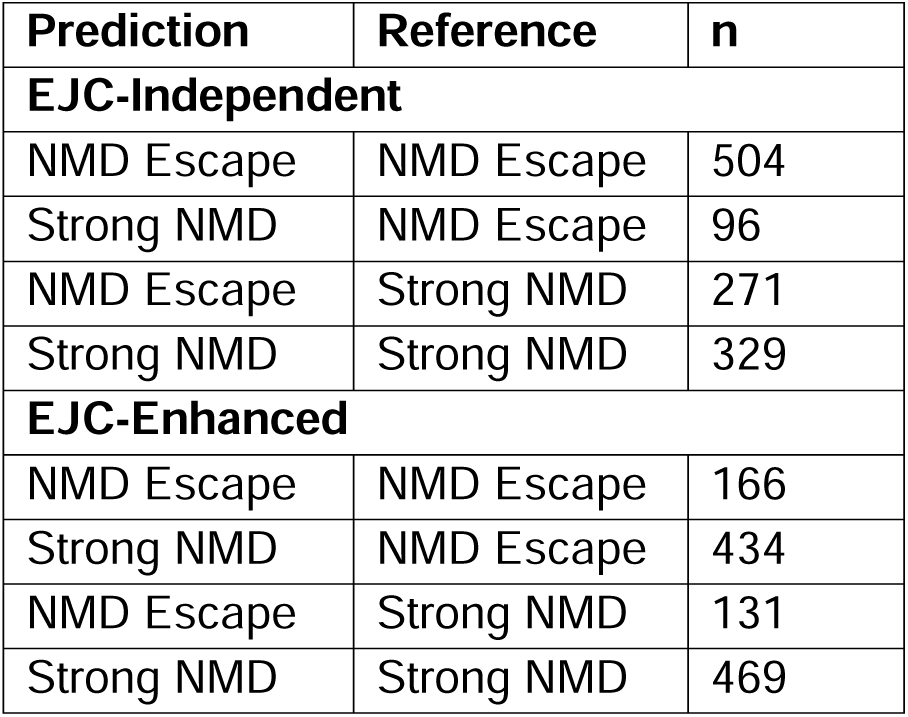
Confusion matrices for the EJC-independent and EJC-enhanced classifier.

| Prediction | Reference | n |
| --- | --- | --- |
| **EJC-Independent** |  |  |
| NMD Escape | NMD Escape | 504 |
| Strong NMD | NMD Escape | 96 |
| NMD Escape | Strong NMD | 271 |
| Strong NMD | Strong NMD | 329 |
| **EJC-Enhanced** |  |  |
| NMD Escape | NMD Escape | 166 |
| Strong NMD | NMD Escape | 434 |
| NMD Escape | Strong NMD | 131 |
| Strong NMD | Strong NMD | 469 |

**Table 2:** Model metrics for the EJC-independent and EJC-enhanced classifier.

|  |  |
| --- | --- |
| **EJC-Independent** |  |
| Sensitivity/ Recall | 0.84 |
| Specificity | 0.55 |
| Pos Pred Value | 0.65 |
| Neg Pred Value | 0.77 |
| Precision | 0.65 |
| F1 | 0.73 |
| Prevalence | 0.50 |
| Detection Rate | 0.42 |
| Detection Prevalence | 0.65 |
| Balanced Accuracy | 0.69 |
| **EJC-Enhanced** |  |
| Sensitivity/ Recall | 0.28 |
| Specificity | 0.78 |
| Pos Pred Value | 0.56 |
| Neg Pred Value | 0.52 |
| Precision | 0.56 |
| F1 | 0.37 |
| Prevalence | 0.50 |
| Detection Rate | 0.14 |
| Detection Prevalence | 0.25 |
| Balanced Accuracy | 0.53 |

## SUPPLEMENTARY DATA

**Supplementary Data Table 1.** DEseq2 results for the EJC-independent library of sequences representing the fold change of DMSO over SMG1i treatment of cells.

**Supplementary Data Table 2.** DEseq2 results for the EJC-enhanced library of sequences representing the fold change of DMSO over SMG1i treatment of cells.

### Declaration of generative AI and AI-assisted technologies in the writing process

During the preparation of this work the authors used Claude.ai to improve language and readability. After using this tool, the authors reviewed and edited the content as needed and take full responsibility for the content of the publication.

